# Modeling and predicting meat yield and growth performance using morphological features of narrow-clawed crayfish with machine learning techniques

**DOI:** 10.1101/2023.03.29.534674

**Authors:** Yasemin Gültepe, Selçuk Berber, Nejdet Gültepe

## Abstract

In this study, predictions of length-weight relationships and meat productivity were generated by machine learning models using measurement data of male and female crayfish in the narrow-clawed crayfish population living in Apolyont Lake. The data set was created using the growth performance and morphometric characters obtained from 1416 crayfish in different years to determine the length-weight relationship and length-meat yield. Statistical methods, artificial intelligence, and machine learning are used due to the difficulty of constructing mathematical models in multi-parameter and multivariate problems. In recent studies, artificial intelligence and machine learning methods give higher accuracy than other prediction methods in large data sets with complex structures. No previous studies have been conducted on such population parameters. The analysis results show that most of the models designed as an alternative to traditional estimation methods in future planning studies in sustainable fisheries, aquaculture, and natural sources management are valid for machine learning and artificial intelligence. Seven different machine learning algorithms were applied to the data set and the length-weight relationships and length-meat yields were evaluated for both male and female individuals. Support Vector Regression (SVR) has achieved the best prediction performance with 0.996 and 0.992 values for the length-weight of males and females, with 0.996 and 0.995 values for the length-meat yield of males and females. The results showed that the SVR outperforms the others for all scenarios regarding the accuracy, sensitivity, and specificity metrics.

## 1 Introduction

Freshwater lobsters, also known as crayfish, which are one of the largest inland forms of decapod crustaceans, which contain economically important species, are represented by 737 species and subspecies in the world [1,2]. The present species in Türkiye is genetically defined as a *Pontastacus leptodactylus* and has different subspecies [2–4]. Crayfish production is done through hunting and breeding in the world. Despite a large number of species, hunting and breeding activities generally focused on the species of only 3 families (Cambaridae, Parastacidae, Astacidae) that are economically important. The amount of crayfish production with hunting was determined as 15,426 tons, excluding China, as of 2015, and Armenia ranks first with a production of 7,380 tons. Considering the production amounts based on species, the most produced crayfish by hunting is *Pontastacus leptodactylus*. Through aquaculture, 787,373 tons of crayfish have been produced in the world. China ranks first with 723,200 tons’ production. *Procambarus clarkii* is the leading species produced by aquaculture with 786,905 tons [5]. In recent years, crayfish plague, overfishing and increased water pollution caused fluctuations in crayfish production. Stock management and alternative production methods should be improved and developed due to fluctuating trends in volumes of crayfish production. For this, first of all, the available data should be used carefully and successfully. In other words, producers have to process data with new technologies and help decision-makers determine the right decisions and strategies for the future. However, it is not possible to manually process and analyze very large amounts of data. There are many different ways in machine learning involving patterns in which relationships will be discovered, each of which determines the type of technique that can be used to make sense of the output from the data. The most commonly used machine learning methods in the literature are; artificial neural networks [6–8], logistic regression [8–9], fuzzy modeling [10], genetic algorithms and programming [11], decision tree [7,9], Bayesian network approach [12–14], random forest [9,15], support vector machine [9,16]. Regression-based modeling techniques are widely used to estimate species distribution. The most commonly used are generalized additive models (GAMs), generalized linear models (GLM), classification and regression trees (CART), and multivariate adaptive regression splines (MARS) [17]. Besides, a modeling approach that integrates a functional network approach with a dynamic Bayesian network model was used to predict trends of different fish and zooplankton species from specific fishing, temperature, and net primary production (Net PP) scenarios [17]. Similarly, Hamilton et al. [13], a Bayesian network approach to developing a habitat suitability model, and Lin et al. [12] were used a Bayesian analysis to account for the combined uncertainty and variability of parameters in the crayfish bioaccumulation model.

For sustainable aquaculture and fisheries, the main focus of this study is to reveal and predict the status of the population by determining the meat yield and growth performance of the narrow-clawed crayfish living in Apolyont Lake in different years and the relationship between length-weight with some morphometric characteristics. Although some studies have estimated the population of some lakes in Turkey, there are no studies that estimated the narrow-clawed crayfish population by using machine learning methods and/or statistical methods in Apolyont Lake. One of the goals of this study is to contribute to the solution to this situation in Apolyont Lake, which is in an important market like Europe. In this context, this study consists of two modules. The first is to apply data preprocessing techniques to all data sets. In the second, Random Forest Regression (RFR), Gradient Boosting Regression (GBR), Decision Tree Regression (DTR), Multilayer Perceptron Regression (MLPR), Support Vector Regression (SVR), Linear Regression (LR) and K-Nearest Neighbors Regression (K-NNR) machine learning methods are executed all together on dataset. For this, various machine learning algorithms were applied to the data set and the best estimation performance of the total length-weight and meat yield of narrow-clawed crayfish was achieved.

## 2 Materials and methods

### 2.1 Study area and data collection

The dataset used in this study is created by using narrow-clawed crayfish data obtained from Apolyont Lake in Balıkesir-Türkiye. Narrow-clawed crayfish were caught totally 1416 sample, which are included 573 (40%) female and 843 (60%) male. Each sample has 22 attributes, and these attributes presented in Table 1. Length measurements of body parts of crayfish are used to determine morphological differences between male and female crayfish among species [15–17]. These measurements are used to determine the comparative growth of populations, the size of crayfish to be put on the market, meat yield and systematic separation. Length measurements of narrow-clawed crayfish broodstock and juvenile individuals were made with a digital caliper with 0.01 mm precision. Length measurements are based on the total length from rostrum point to telson point. In the measurement of the weight of narrow-clawed crayfish, the weight of the broodstock was measured with a 0.01 g precision weighing, while the weight of the juvenile individuals was measured with a 0.0001 g precision scale [18].

**Table 1.**
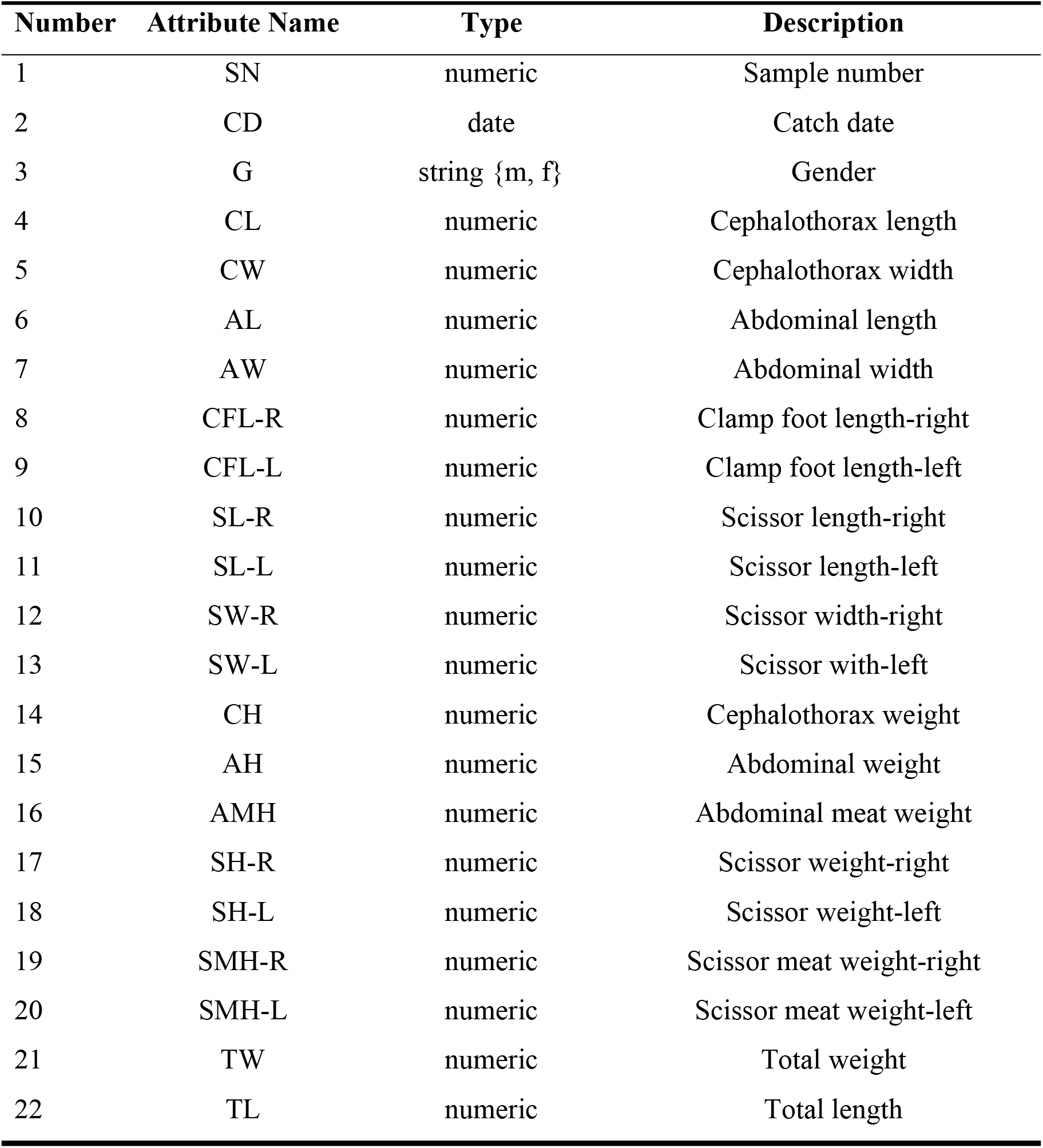
Attribute information.

### 2.2 Length-weight relationship and meat yield

Regression analysis is generally used to determine the relationship between body length and weight of crustaceans [19]. As in fish, there is a nonlinear relationship between length and weight in the form of Equation 1 in crayfish. If the logarithms of both sides are taken in this equation, the length-weight relationship becomes linear as Equation 2 [19,20].

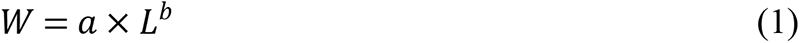

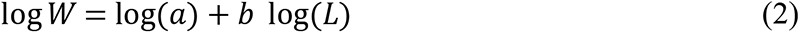

Abbreviations: L, total length (TL); W, total weight (TW); a and b, constant parameters of the equation

The length-weight relationship of narrow-clawed crayfish was investigated in terms of total length (TL) – total weight (TW) relationship. Accordingly, regression equations, curves and correlation coefficients were calculated. While checking the significance of the calculated b value, the test statistic value was calculated using Equation 3.

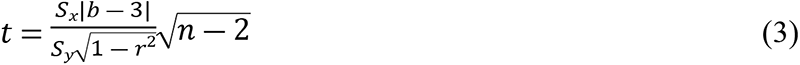

Abbreviations: Sx, standard deviation of log (L) values; Sy, standard deviation of log (W) values); n, the number of individuals used in the calculation; r^2^, coefficient of determination of the log (L) and log (W) values

To determine the meat yield, the abdomen, claws and scissor were cut with the help of a scalpel, and the meats inside were directly weighed and their weights were determined [21].

### 2.3 Machine learning techniques

Machine learning is the general name of computer algorithms that can learn the solution of the problem, which is addressed with complex pattern detection and data-based decision-making features [22]. Classification is distributing data to classes in a data set according to their attributes. Classification algorithms analyze the relationships between class labels and other features in a given training set [23]. The success of the model has decided which class belongs to the new item and this is tested with the help of model. In this study, the modeling and prediction of population growth of crayfish in Lake Apolyont were made using popular machine learning algorithms Random Forest Regression (RFR), Gradient Boosting Regression (GBR), Decision Tree Regression (DTR), Multilayer Perceptron Regression (MLPR), Support Vector Regression (SVR), Linear Regression (LR) and K-Nearest Neighbors Regression (K-NNR). Their achievements have been compared with each other.

RFR is a collection of decision trees, each independently of the other and based on a random sample of training data using the same distribution. This method creates many decision trees during the training, and then, during the estimation, the classification of these decision trees is used, and the class of the input is decided by a majority vote. RF regression is a regression model that uses more than one decision tree to produce more compatible models and make accurate predictions. It is discrete because it uses decision trees. That is, it produces the same results for the desired predictions in a certain range [9,23,24].

GBR is a machine learning algorithm and a model developed to improve the prediction of decision trees. It is an algorithm developed by Friedman [25]. GBR algorithm is similar to other decision trees, it produces a stronger ring by making errors from each other’s weaknesses. According to the GBR algorithm, a prediction function is first created in the first iteration and these functions are called trees. While creating the next tree, the error rates of the trees created before are kept in memory. The difference between the estimates and the observations is calculated and a loss function is obtained from these differences. In the second iteration, the difference between the predictions and the observations is calculated by combining the prediction and loss functions. Thus, it is tried to increase the success of the estimation function by adding it continuously and it is ensured that the error rate approaches zero.

DTR observes the properties of an object and trains a model in the structure of a tree to predict future data to produce meaningful continuous output. Continuous output means that the output/result is not discrete, i.e. only represented by a discrete, known set of numbers or values [23,26].

MLPR is a neural network with one or more hidden layers between the input layer and the output layer. MLPR can classify nonlinear data through several hidden layers and nonlinear enable functions such as ReLU and tanh. In particular, regression analysis using MLPR does not require the assumption of a statistical relationship between independent and dependent variables. For this reason, MLPR is widely used as an algorithm for regression in various fields [27–29].

SVR is a support vector machine (SVM) implementation that generates an actual number as output. SVM can be applied to regression problems by importing an alternative loss function. SVR is built on the principle of inherent risk minimization to solve complex problems [30,31].

LR is a supervised learning algorithm that uses a parametric model and a linear approach for a prediction problem [9,23,32].

K-NNR for machine learning is well known to have been introduced as a non-parametric approach used to classify fields and perform regressions. Within these two areas, the input data contains the closest training examples in the feature area. In K-NNR, the output is the property value of the object. This value is the average of the values of the k-nearest neighbors [31,33].

Data preprocessing, which will increase the quality of raw data to be used in the study, is one of the most important processes that have a direct positive effect on computer science and the performance of all algorithms. Since machine learning algorithms are generally data-driven structures, various operations such as cleaning, scaling, reducing, and normalization have significant effects on prediction accuracy [34,35]. In this study, after the raw data were arranged and determined, they were subjected to a normalization process with min-max normalization. The applied normalization formula is given in Equation 4.

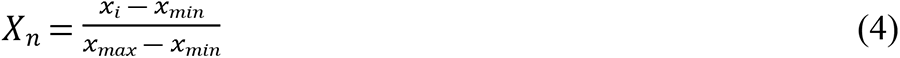

In this formula, each input (x_i_) value is linearly normalized (x_n_) between 0 and 1 by finding the minimum (x_min_) and maximum (x_max_) values of the raw data set [23,31]. Of the available data, 70% was used for training the machine learning model and 30% for testing it. The flow chart of the proposed method in this study is given in Fig. 1.

**Fig 1.**
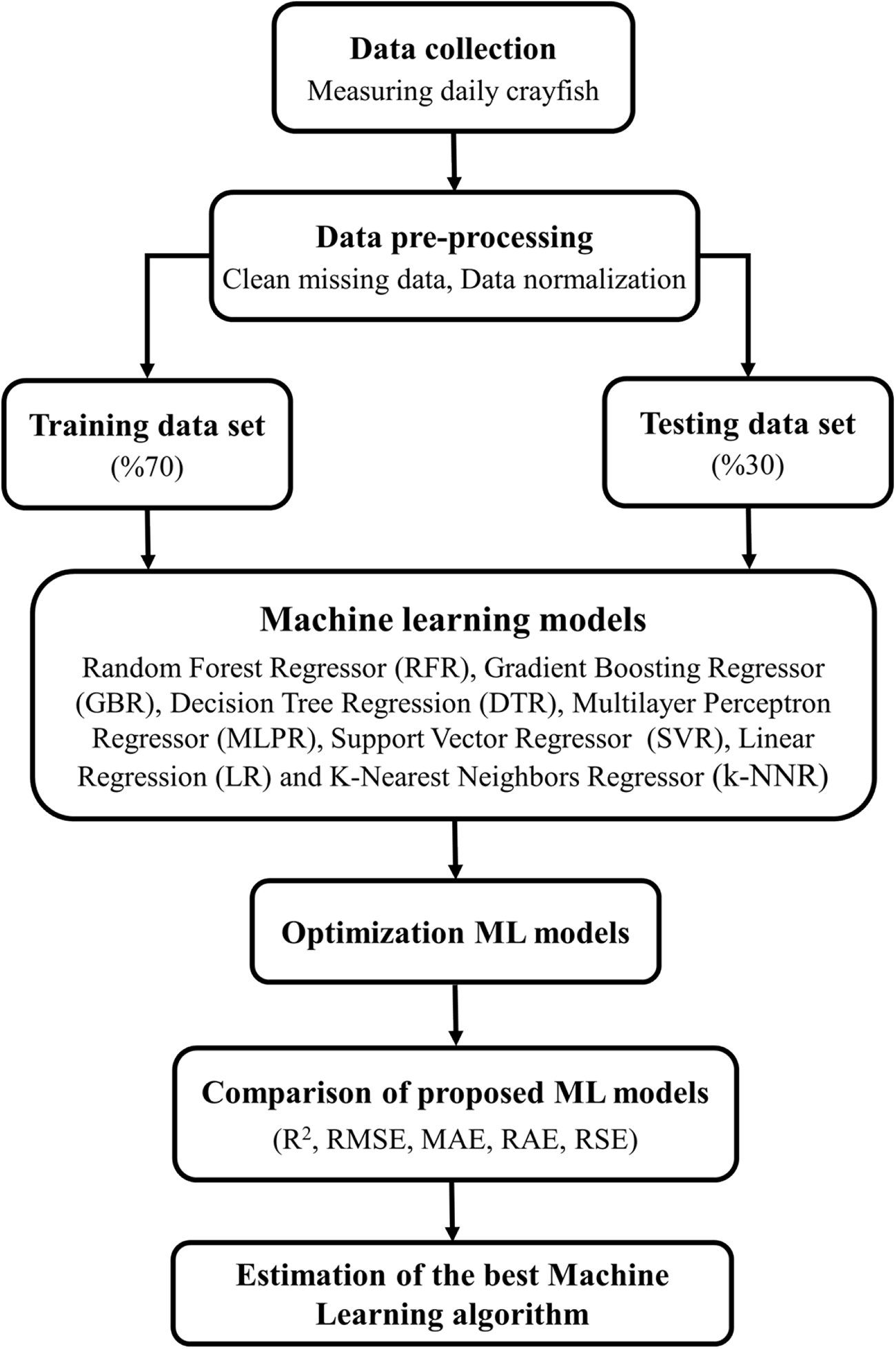
Flow diagram of the system. (Abbreviations: R^2^, determination coefficient; RMSE, root mean square error; MAE, mean absolute error; RAE, relative absolute error; RSE, root relative square error)

The models were made using a desktop on the operating system Windows 10 Pro operating system with the following hardware configuration: Intel(R) Core(TM) i7-7700HQ, 2.80 GHz processor speed, and 16.0 GB of RAM. Python language was used in the present study.

### 2.4 Performance metrics

R-squared (determination coefficient) (R^2^), root mean square error (RMSE), mean absolute error (MAE), relative absolute error (RAE), and root relative square error (RSE) values were used to measure the estimation performance of the models in the study. Using these metrics, it can be decided which technique is most suitable for this data set. In linear regression, R^2^ (Equation 5) is a measure of how close the data points are to the fit line. It is also known as the coefficient of determination. The R^2^ is a metric that represents prediction performance for regression models. It is a positive value between 0 and 1. The ideal value for R^2^ is 1. The closer the R^2^ value is to 1, the better the model. RMSE (Equation 6), prediction errors, is a measure of how far the regression line is from the data points. MAE (Equation 7) is the error rate of the growth forecast model. RAE (Equation 8) takes the total absolute error and normalizes it by dividing it by the simple estimator’s total absolute error. RSE (Equation 9) gives the square root of the sum of the squares of the differences between the estimated value and the true value to the sum of the squares of the differences between the true values and the mean value [36]. The lower the calculated values for the four error metrics and the closer the determination coefficient is to 1, the more accurate the results are. These 5 different metrics are not sufficient wit alone in terms of the calculated values but are meaningful when evaluated together.

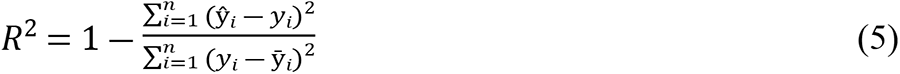

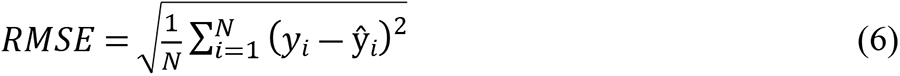

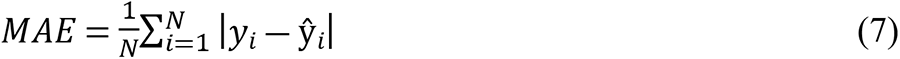

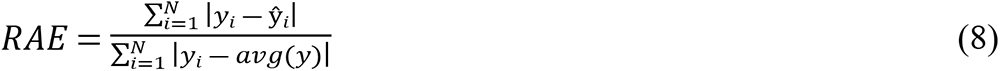

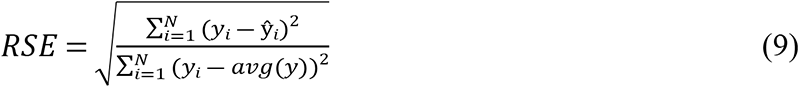

## 3 Results

Length-weight relationships and meat yields are used to compare the characteristics of different crayfish populations. In this study, the weights of crayfish according to gender are associated with their lengths. It was seen that the crayfish ranged between 23-71 mm carapace lengths. The carapace length has measured a minimum of 23 maximum of 70 mm in females, a minimum of 28 maximum of 71 mm in males, and an average of 44 mm in all individuals. When the weight distributions of the crayfish were examined, it was determined that the live weight was between 2.5-92.4 g. The weight of female individuals ranged between 2.5-72.4 g, and the weight of male individuals ranged between 2.5-92.4 g. The average total weight measured was 20.0 g in females, and 21.9 g in males. Although there was a linear relationship between meat amounts and carapace lengths in male and female crayfish, it was determined that this relationship was stronger in male crayfish. Total meat yield was calculated as 16.45% on average in the examinations performed on a total of 1416 individuals.

Seven different machine learning algorithms were run on the training data and the total weight performance metric results according to the length measurements of male individuals are given in Table 2 and the results of female individuals are given in Table 3. Likewise, the abdominal meat yield performance metric results according to the length measurements of male and female individuals are given in Table 4 and Table 5, respectively. The best length-weight and length-meat yield performance metric results for all individuals were found with SVR. According to the SVR, accuracy levels were found in the length-weight prediction results of males (R^2^=0.996; RMSE=0.979) and females (R^2^=0.992; RMSE=0.966), and the length-meat yield prediction results of males (R^2^=0.996; RMSE=0.098) and females (R^2^=0.995; RMSE=0.977) were 99.6%, 99.2%, 99.6%, and 99.5%, respectively. The lowest accuracy levels were found with MLPR for female length-weight, male and female length-meat yield performance metric results. On the other hand, in male length-weight performance metric results, the lowest accuracy level was found with DTR.

**Table 2.**
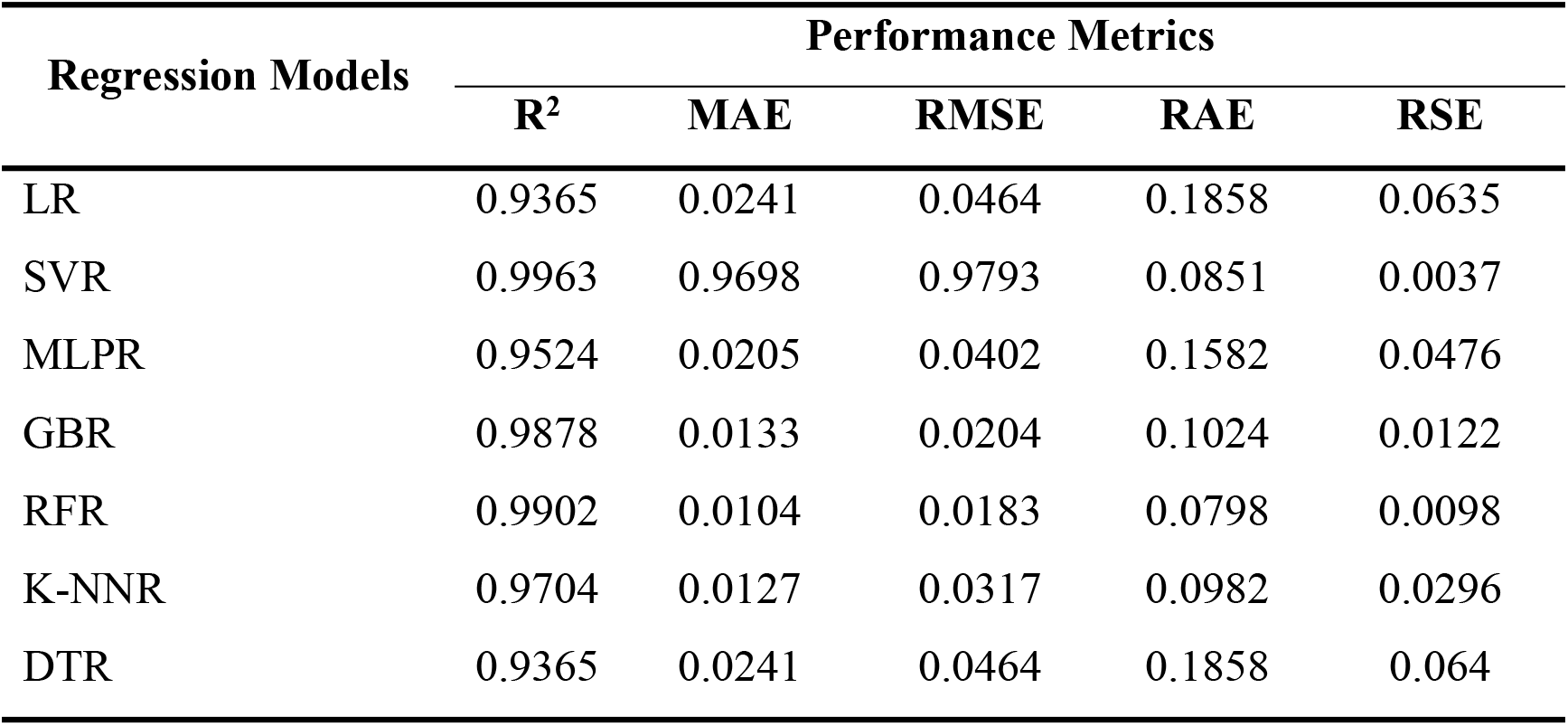
Performance metric results for total weight according to length measurement values in male crayfish. (Abbreviations: R^2^, determination coefficient; RMSE, root mean square error; MAE, mean absolute error; RAE, relative absolute error; RSE, root relative square error; LR, Linear Regression; SVR, Support Vector Regression; MLPR, Multilayer Perceptron Regression; GBR, Gradient Boosting Regression; RFR, Random Forest Regression; K-NNR, K-Nearest Neighbors Regression; DTR, Decision Tree Regression)

| Regression Models | Performance Metrics |  |  |  |  |
| --- | --- | --- | --- | --- | --- |
| | $R^2$ | MAE | RMSE | RAE | RSE |
| LR | 0.9365 | 0.0241 | 0.0464 | 0.1858 | 0.0635 |
| SVR | 0.9963 | 0.9698 | 0.9793 | 0.0851 | 0.0037 |
| MLPR | 0.9524 | 0.0205 | 0.0402 | 0.1582 | 0.0476 |
| GBR | 0.9878 | 0.0133 | 0.0204 | 0.1024 | 0.0122 |
| RFR | 0.9902 | 0.0104 | 0.0183 | 0.0798 | 0.0098 |
| K-NNR | 0.9704 | 0.0127 | 0.0317 | 0.0982 | 0.0296 |
| DTR | 0.9365 | 0.0241 | 0.0464 | 0.1858 | 0.064 |

**Table 3.** Performance metric results for total weight according to length measurement values in female crayfish. (Abbreviations: R^2^, determination coefficient; RMSE, root mean square error; MAE, mean absolute error; RAE, relative absolute error; RSE, root relative square error; LR, Linear Regression; SVR, Support Vector Regression; MLPR, Multilayer Perceptron Regression; GBR, Gradient Boosting Regression; RFR, Random Forest Regression; K-NNR, K-Nearest Neighbors Regression; DTR, Decision Tree Regression)

| Regression Models | Performance Metrics |  |  |  |  |
| --- | --- | --- | --- | --- | --- |
|  | R<sup>2</sup> | MAE | RMSE | RAE | RSE |
| LR | 0.9429 | 0.0212 | 0.0295 | 0.2234 | 0.057 |
| SVR | 0.992 | 0.9533 | 0.9661 | 0.1144 | 0.008 |
| MLPR | 0.9441 | 0.0201 | 0.0291 | 0.2135 | 0.0559 |
| GBR | 0.9899 | 0.0094 | 0.0123 | 0.0992 | 0.01 |
| RFR | 0.9895 | 0.0088 | 0.0126 | 0.0928 | 0.0104 |
| K-NNR | 0.9798 | 0.0104 | 0.0175 | 0.1094 | 0.0202 |
| DTR | 0.9342 | 0.0198 | 0.0316 | 0.2089 | 0.0658 |

**Table 4.** Total length and abdominal meat yield efficiency of male crayfish. (Abbreviations: R^2^, determination coefficient; RMSE, root mean square error; MAE, mean absolute error; RAE, relative absolute error; RSE, root relative square error; LR, Linear Regression; SVR, Support Vector Regression; MLPR, Multilayer Perceptron Regression; GBR, Gradient Boosting Regression; RFR, Random Forest Regression; K-NNR, K-Nearest Neighbors Regression; DTR, Decision Tree Regression)

| Regression Models | Performance Metrics |  |  |  |  |
| --- | --- | --- | --- | --- | --- |
|  | R<sup>2</sup> | MAE | RMSE | RAE | RSE |
| LR | 0.9347 | 0.0361 | 0.0486 | 0.2470 | 0.0653 |
| SVR | 0.9326 | 0.0386 | 0.0494 | 0.2641 | 0.0675 |
| MLPR | 0.9186 | 0.0384 | 0.0543 | 0.2631 | 0.0814 |
| GBR | 0.9910 | 0.0138 | 0.0180 | 0.0942 | 0.009 |
| RFR | 0.9929 | 0.0099 | 0.0161 | 0.0674 | 0.0071 |
| K-NNR | 0.9744 | 0.0201 | 0.0304 | 0.1375 | 0.0256 |
| DTR | 0.9251 | 0.0318 | 0.0521 | 0.2178 | 0.0749 |

**Table 5.** Total length and abdominal meat yield efficiency of female crayfish. (Abbreviations: R^2^, determination coefficient; RMSE, root mean square error; MAE, mean absolute error; RAE, relative absolute error; RSE, root relative square error; LR, Linear Regression; SVR, Support Vector Regression; MLPR, Multilayer Perceptron Regression; GBR, Gradient Boosting Regression; RFR, Random Forest Regression; K-NNR, K-Nearest Neighbors Regression; DTR, Decision Tree Regression)

| Regression Models | Performance Metrics |  |  |  |  |
| --- | --- | --- | --- | --- | --- |
| | $R^2$ | MAE | RMSE | RAE | RSE |
| LR | 0.9251 | 0.0318 | 0.0521 | 0.2177 | 0.0749 |
| SVR | 0.8901 | 0.0467 | 0.055 | 0.3637 | 0.1099 |
| MLPR | 0.8878 | 0.0420 | 0.0556 | 0.3266 | 0.1122 |
| GBR | 0.9859 | 0.0146 | 0.0197 | 0.1135 | 0.0141 |
| RFR | 0.9879 | 0.0123 | 0.0183 | 0.0956 | 0.0121 |
| K-NNR | 0.9519 | 0.0255 | 0.0364 | 0.1985 | 0.0481 |
| DTR | 0.9015 | 0.0372 | 0.0521 | 0.2893 | 0.0986 |

## 4 Discussion

The exact determination of the population size is possible by hunting all individuals. Since it is not possible to do this in practice, the population size is tried to be determined with the help of the population parameters explained here. Because, knowing the current status of the population is important in determining the fishing strategies in fisheries management. The ideal is to make estimates of population size at regular intervals. However, this is a tiring job that requires money and effort. The change in the population is observed by doing it at least every few years. Because population size depends on time in a certain period, death, reproduction, migration, growth, hunting, abundance of food, predisposing creatures, etc. changes in number and weight with the effect of factors [37,38]. Stock monitoring studies are studies that need to be carried out uninterruptedly for the sustainability of populations and determination of growth characteristics in population parameters is also an important parameter. Therefore, it was aimed to determine these growth characteristics of crayfish in this study.

Machine learning provides a neutral approach to recognizing unknown interactions and deriving predictions that have the potential to aid in meaningful feature selection. The correlation values produced as a result of the algorithm estimates show the correlation between length-weight and length-meat yield measurement values and length-weight and length-meat yield prediction values. Since this study is based on the properties that are effective in the correct estimation of the length-weight and length-meat yield values with the determination of the correct length-weight and length-meat yield prediction, they are evaluated on the positive correlation values formed as a result of the estimates of the algorithms. Because the proximity of correlation values to +1 indicates the closeness of length-weight and length-meat yield estimates to length-weight and length-meat yield measurements.

Some of the growth parameters of Tigris loach (*Oxynoemacheilus tigris*) were estimated by using both length-weight relationship and artificial neural network (ANN) between 2014-2015 from 14 different stations in Karasu and the two methods were compared with each other. It has been observed that there is a high affinity between the measured and predicted data and the values obtained with ANN are closer to the real values [39]. Also Ozcan [40] pointed out that ANN can be used as an alternative method for estimation of population. In the present study, SVR performance metric results are 0.996 and 0.992 values both for the length-weight of male and female individuals, with 0.996 and 0.995 values and for the length-meat yield of males and females are closer to the values of measurement. SVR stands out as the method with the best R2 and error values. SVR is both linear and non-linear [31] due to this feature, it is considered that the SVR provides better performance in evaluating the population structure. Similarly, Benzer et al. [8], and Benzer & Benzer [41], showed that ANN could be a superior estimation tool compared to the length-weight for the growth predictions of narrow-clawed crayfish in Hirfanlı Dam Lake and Uluabat Lake, respectively. In addition, SVM gave 80% accuracy in classifying sea bream feeding trials according to hematological blood parameters. In addition, SVM provided 80% accuracy in classifying sea bream (*Sparus aurata*) feeding trials according to hematological blood parameters [9], but K-NN gave better results than SVM in *Alburnus tarichi* population analysis [31]. All these studies showed that artificial intelligence applications such as ANN and machine learning etc. should be specific to data. Before evaluating a population in the long term, the first thing to do is to determine which application is appropriate for that population. The results of this study showed that SVR is the most successful application to evaluate the crayfish population living in Apolyont Lake. To better evaluate the results, the prediction scores and actual values were given for male length-weight in Figure 2, for female length-weight in Figure 3, for male length-meat yield in Figure 4, and for female length-meat yield in Figure 5.

**Fig 2.**
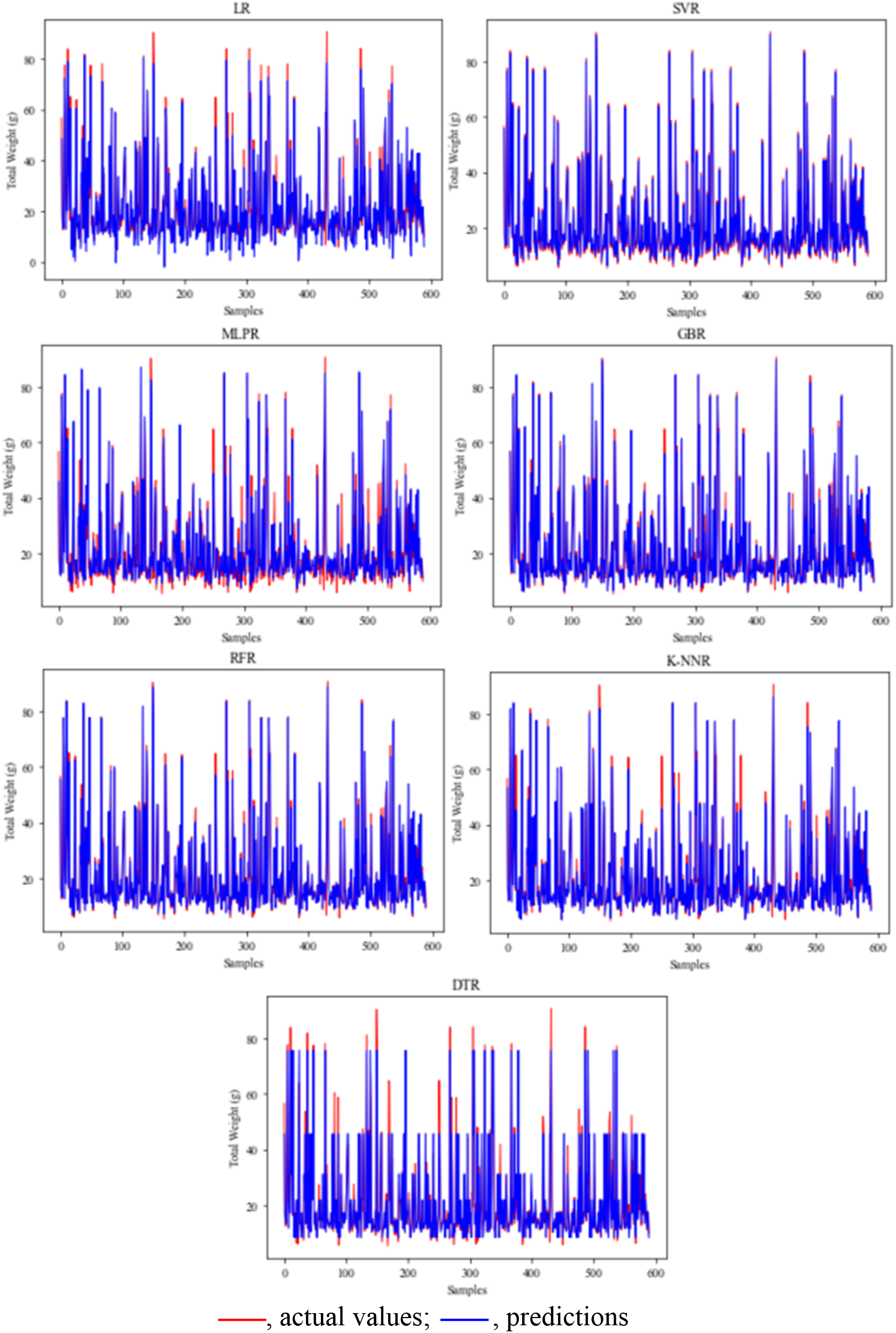
Length-weight prediction and actual values in male crayfish. (Abbreviations: LR, Linear Regression; SVR, Support Vector Regression; MLPR, Multilayer Perceptron Regression; GBR, Gradient Boosting Regression; RFR, Random Forest Regression; K-NNR, K-Nearest Neighbors Regression; DTR, Decision Tree Regression)

**Fig 3.**
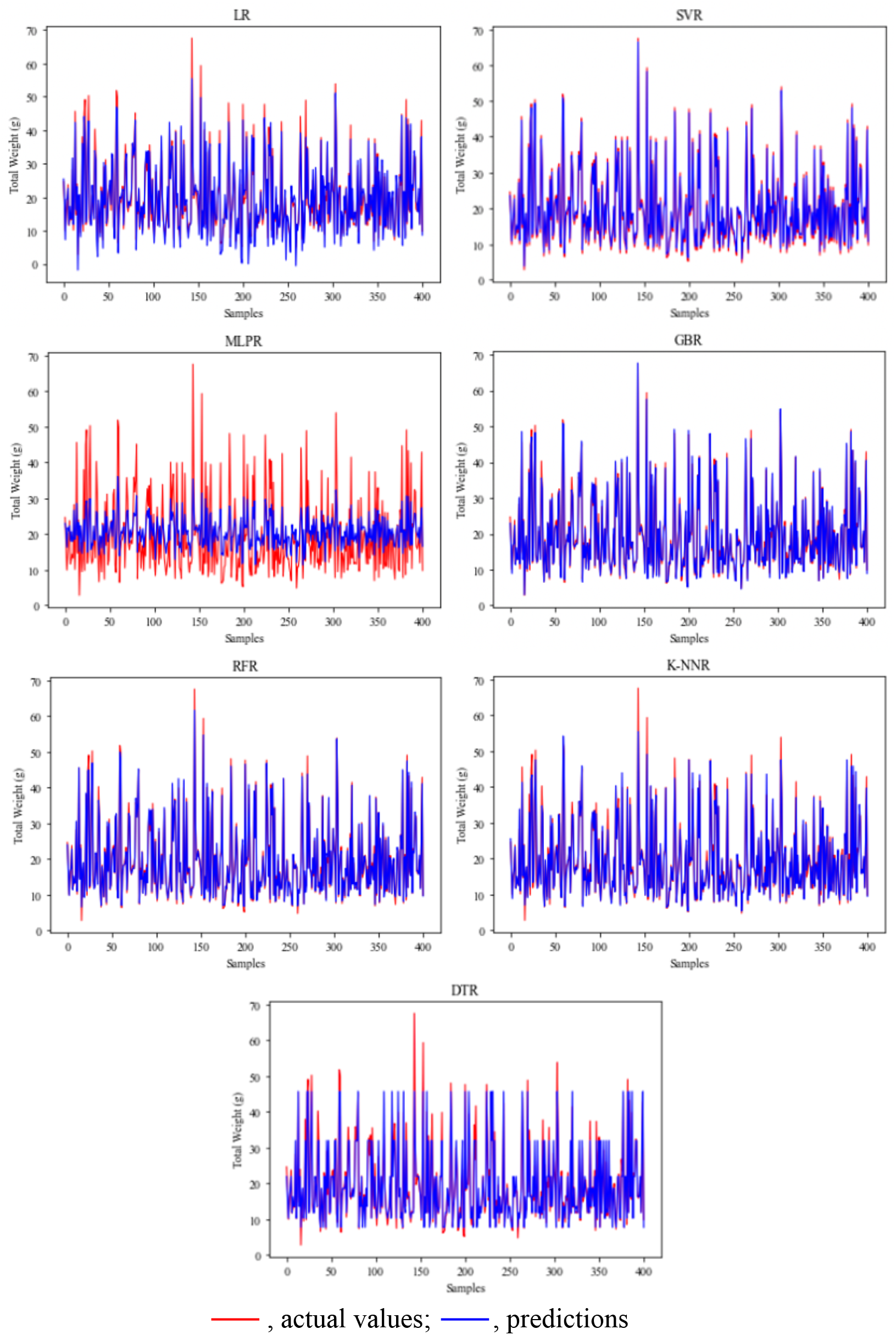
Length-weight prediction and actual values in female crayfish. (Abbreviations: LR, Linear Regression; SVR, Support Vector Regression; MLPR, Multilayer Perceptron Regression; GBR, Gradient Boosting Regression; RFR, Random Forest Regression; K-NNR, K-Nearest Neighbors Regression; DTR, Decision Tree Regression)

**Fig 4.**
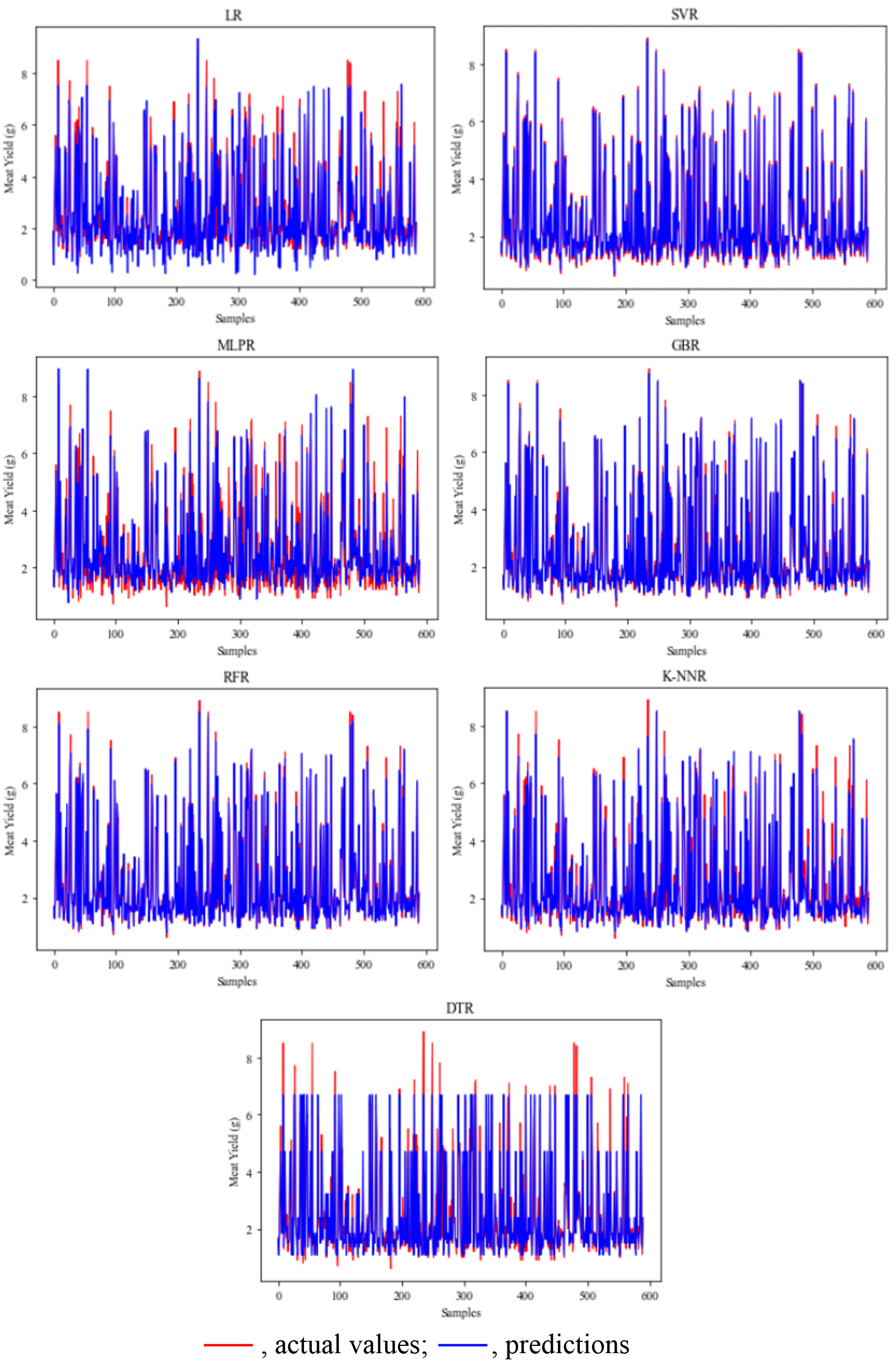
Meat yield prediction and actual values in male crayfish. (Abbreviations: LR, Linear Regression; SVR, Support Vector Regression; MLPR, Multilayer Perceptron Regression; GBR, Gradient Boosting Regression; RFR, Random Forest Regression; K-NNR, K-Nearest Neighbors Regression; DTR, Decision Tree Regression)

**Fig 5.**
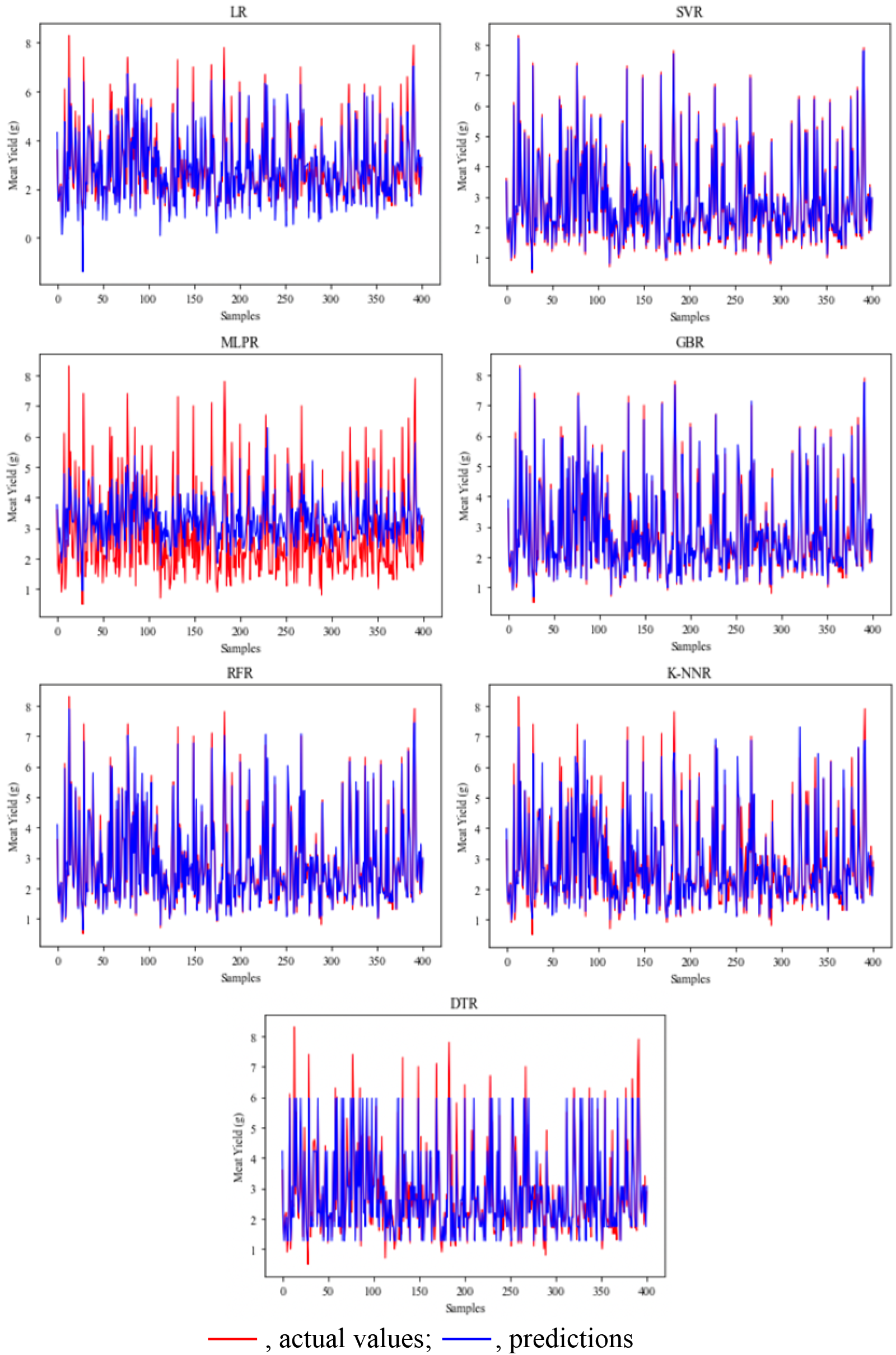
Meat yield prediction and actual values in female crayfish. (Abbreviations: LR, Linear Regression; SVR, Support Vector Regression; MLPR, Multilayer Perceptron Regression; GBR, Gradient Boosting Regression; RFR, Random Forest Regression; K-NNR, K-Nearest Neighbors Regression; DTR, Decision Tree Regression)

Ecological factors determining the presence of white-clawed crayfish (*Austropotamobius pallipes*) were evaluated using SVM, a machine learning method. It was determined that the models without feature selection, the models created by applying Goldberg’s genetic algorithm after the feature selection, and the models built after selecting inputs using the four supervised-filter evaluators had a classification accuracy degree of 70.84%, 73.92%, 76.62%, respectively [16]. Applied model in this study have higher accuracy as well as found closer to the real values. Thus, the study results of Zelaya [34] were supported and the accuracy of the results obtained from this study was observed. Besides, Tirelli et al. [7] used logistic regression, decision tree models, and artificial neural network to manage data on the presence/absence of native white-clawed crayfish. They obtained better performance from logistic regression and decision tree models using the artificial neural network model.

However, a hybrid three-dimensional (3D) dissolved oxygen content estimation model based on a radial basis function neural network, K-means, and subtractive clustering effectively demonstrated the three-dimensional distribution in predicting changes in dissolved oxygen content on crap pools [42]. Many crayfish cannot be sold at the end of the sales period due to errors made in production planning, growth estimation, transportation, grading processes, and the application of the appropriate stock density. This situation causes a loss of resources, energy, and capital in both production and sales stages. In present study, it was seen that due to the multivariate nature of predicting the growth rate of crayfish populations, instead of creating hybrid models, it can be done by using machine learning methods that give faster and more accurate results in complex structured data sets.

## 5 Conclusion

This is the first study to use machine learning methods to predict the total length-total weight and total length-meat yield performance of the narrow-clawed crayfish population of Apolyont Lake. The results show that machine learning methods have the ability to predict population growth rates and size in fisheries and aquatic populations. Among these seven machine learning applications implemented, SVR gave the best performance in terms of both R^2^ value and error metrics. In the study, it was seen that the values obtained by machine learning have high performance and are closer to the real values. The experimental results of the study show that the proposed method can be used as an effective population measurement estimation tool. Moreover, this article, which presents a series of empirical analyzes with the results discussed, is considered to be valuable for professionals of natural resource management, fisheries, aquaculture, and their sustainability.

## Acknowledgement

This research received no external funding. The data that support the findings of this study are available upon request from the authors.

